# The FH2 domain of formin proteins is critical for platelet cytoskeletal dynamics

**DOI:** 10.1101/589861

**Authors:** Hannah L.H. Green, Malou Zuidscherwoude, Steven G. Thomas

## Abstract

Reorganisation of the actin cytoskeleton is required for proper functioning of platelets following activation in response to vascular damage. Formins are a family of proteins which regulate actin polymerisation and cytoskeletal organisation. Several formin protein are expressed in platelets and so we used an inhibitor of formin mediated actin polymerisation (SMIFH2) to uncover the role of these proteins in platelet spreading. Pre-treatment with SMIFH2 completely blocks platelet spreading in both mouse and human platelets through effects on the organisation and dynamics of actin and microtubules. However, platelet aggregation and secretion are unaffected. SMIFH2 also caused a decrease in resting platelet size and disrupted the balance of tubulin post-translational modification. These data therefore demonstrated an important role for formin mediated actin polymerisation in platelet spreading and highlighted their importance in cross talk between the actin and tubulin cytoskeletons.

**Key Points:**

- Inhibition of FH2 domains blocks platelet spreading and disrupts actin and microtubule organisation
- Inhibition of FH2 domains causes a reduction in resting platelet size but not by microtubule coil depolymerisation
- FH2 domains play a role in the post-translational modification of microtubules

**Visual abstract:** 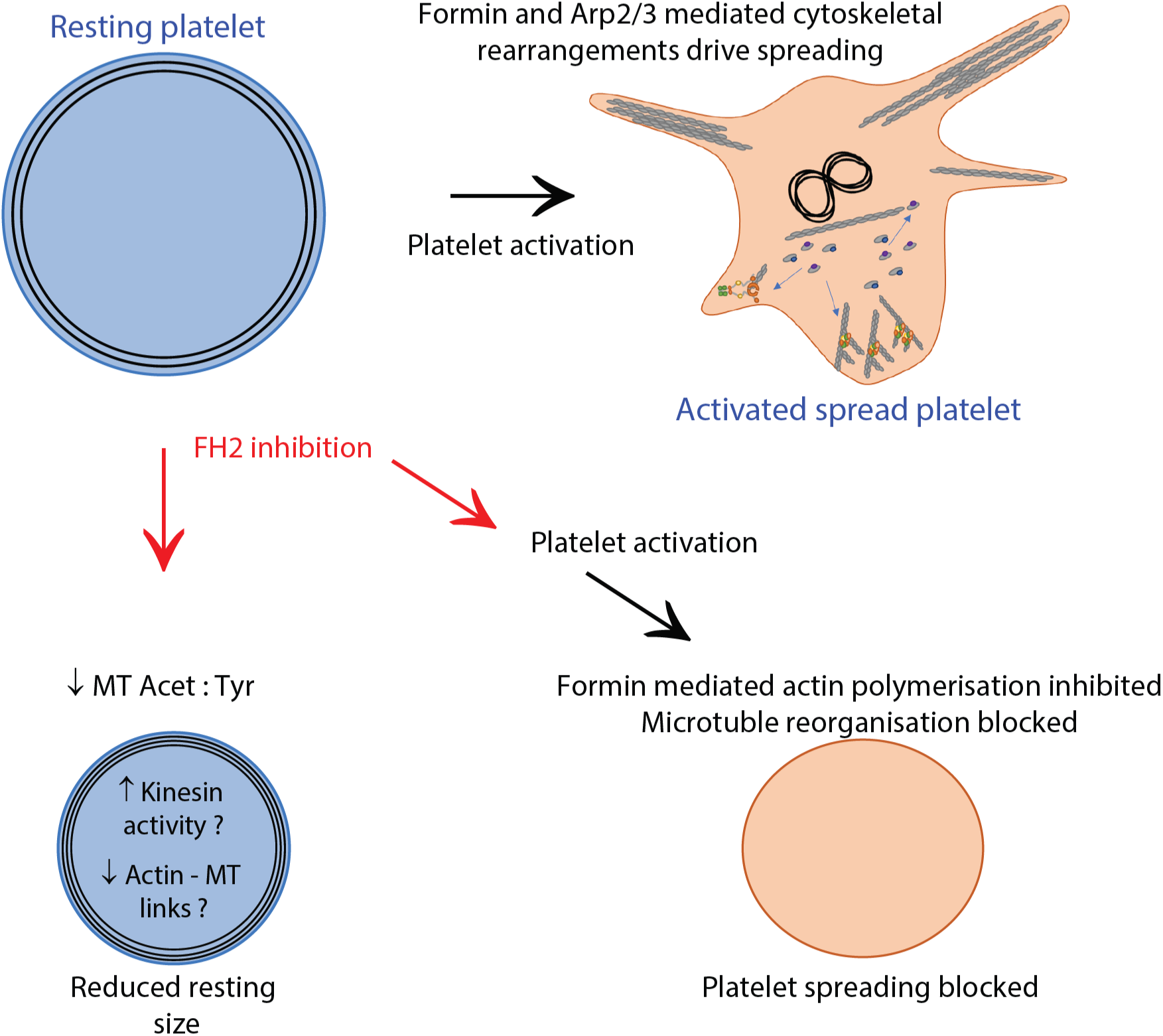

## Introduction

Platelets require a dynamic actin and tubulin cytoskeleton for proper maintenance of their resting size and shape, and for their response to vascular damage. Formins are a family of 15 actin nucleating factors that promote the assembly of linear actin filaments downstream of the Rho family of small GTPases ^[1]^. Formin proteins contain a number of domains including FH1 and FH2 domains ^[1]^ and are known to functioning as homodimers ^[2]^. Profilin bound actin monomers associate with the FH1 domain of the formin dimer and are added to the growing actin filament by the action of FH2 domains, which act to both elongate the filament and protect it from capping proteins ^[3, 4, 5]^. In addition to their role in nucleating actin filaments, formins have been shown to play an important role in microtubule organisation and dynamics, including alignment of actin and microtubules, microtubule stabilisation and microtubule bundling ^[6, 7]^ indicating a role for formin proteins in coordination of actin and microtubule cytoskeletons in cells.

Zuidscherwoude *et al.* ^[7]^ have reviewed expression databases for the 15 formin proteins in developing megakaryocytes and platelets at both the DNA and protein level. We reported that only 4 of the 15 formins were expressed in human or mouse platelets; these being DAAM1, mDia1, FHOD1 & INF2. The presence of DAAM1, mDia1 and FHOD1 has been confirmed in platelets by western blotting ^[8]^. Studies on the mDia1 knockout mouse indicated no major phenotype in terms of platelet activation, aggregation or spreading, possibly due to functional redundancy between the expressed formin members ^[8]^. Despite this, mDia1 has been shown to play a role in megakaryocyte development and proplatelet formation (PPF) as knockdown of mDia1 using shRNA resulted in increased PPF ^[9]^. Further studies on gain of function mutations in mDia1 have shown that constitutively active mDia1 leads to reduced PPF, and patients expressing gain of function mutations display (macro)thrombocytopenia ^[9, 10, 11]^. Despite these studies, the contribution of formin proteins to platelet actin and microtubule organisation remains unclear.

SMIFH2 was identified in 2008 by screening small molecules for their ability to inhibit actin assembly by mDia1 and mDia2 ^[12]^. This activity was attributed to the FH2 domain, as the ability to inhibit actin polymerisation persisted when profilin and the FH1 domain were absent. In mammalian fibroblasts, SMIFH2 inhibited formin dependent migration and reduced membrane integrity, but had little effect on Arp2/3 complex-dependent structures such as lamellipodia ^[12]^. However, the effects of FH2 domain inhibition on the platelet cytoskeleton are not defined. Therefore, in the present study, human and mouse platelets were visualised with fluorescence microscopy to observe changes to morphology, and actin and tubulin dynamics in response to SMIFH2 treatment. Assays were performed with resting platelets and platelets undergoing spreading in response to activation by fibrinogen. Furthermore, the stability of microtubules following SMIFH2 treatment was investigated by fluorescence microscopy and western blotting to detect post-translationally modified populations of α-tubulin in resting platelets, indicative of stable and dynamic forms of the protein.

## Methods

### Mice

C57Blk6 or Lifeact-GFP ^[13]^ mice were maintained in IVCs under 12 h light/dark cycle at a constant temperature of 20 °C with food and water given *ad libitum* at BMSU, University of Birmingham, UK. All experiments were performed in accordance with UK laws (Animal [Scientific Procedures] Act 1986) with approval of local ethics committee (Birmingham Animal Welfare and Ethical Review Board – AWERB) under a Home Office approved project licence.

### Platelet preparation

Human washed platelets were prepared from blood samples donated by healthy, consenting volunteers (local ethical review no: ERN-11-0175). Blood was drawn via venipuncture into sodium citrate as the anticoagulant and then acid/citrate/dextrose (ACD) was added to 10% (v:v). Blood was centrifuged at 200 × g for 20 min. Platelet rich plasma (PRP) was obtained and then centrifuged at 1000 × g for 10 min in the presence of 0.1 *µ*g ml^-1^ prostacyclin (Cayman Chemicals). Plasma was removed and the platelet pellet was resuspended in modified Tyrodes buffer (134mM NaCl, 0.34mM Na_2_HPO_4_, 2.9mM KCl, 12mM NaHCO_3_, 20mM HEPES, 5mM glucose, 1mM MgCl_2_; pH 7.3) containing ACD and 0.1 *µ*g ml^-1^ prostacyclin before being centrifuged for 10 min at 1000 × g. The washed platelet pellet was resuspended in modified Tyrodes buffer, left to rest for 30 min and the platelet count determined using a Coulter Counter (Beckman Coulter). Platelet count was adjusted as required using modified Tyrodes buffer.

Mouse washed platelets were prepared from blood drawn from the vena cava of CO_2_ narcosed mice directly into 100 *µ*l ACD. PRP was obtained by centrifugation at 200 × g for 6 min. Washed platelets were prepared via centrifugation of PRP at 1000 × g in the presence of prostacyclin (0.1 *µ*g^-1^) for 6 min. The pellet was resuspended in modified Tyrodes buffer and counted as per human platelets.

### Platelet spreading

For fixed cell immunofluorescence imaging, platelets were diluted to 2 × 10^7^ ml^-1^ in modified Tyrodes buffer and allowed to spread on fibrinogen (Enzyme Research) coated coverslips (100 *µ*g ml^-1^) for 45 mins at 37 °C, 5 % CO_2_. Where indicated, platelets spread on fibrinogen had 0.1 U ml^-1^ thrombin (Sigma) added just prior to adding the platelets to the coverslip. For inhibitor studies, SMIFH2 (Merck Millipore) was added to the cells at the required concentration for 10 mins at room temperature prior to spreading. At the end of the incubation, non adhered platelets were removed by washing once in PBS and then fixed for 10 mins in 10 % neutral buffered formalin (Sigma-Aldrich).

For live cell imaging experiments, washed human or Lifeact-GFP mouse platelets were prepared as above and diluted to 2 × 10^8^ ml^-1^ before being incubated with 1 *µ*M SiR-tubulin (Spirochrome) at 37 °C for 1 hour. Platelets were diluted to 4 × 10^7^ ml^-1^ and incubated with SMIFH2 at the required concentration before adding to fibrinogen coated glass bottomed dishes (MatTek) and imaging.

For resting platelet experiments, washed human and mouse platelets at 4 × 10^7^ ml^-1^ were incubated with SMIFH2 at 37 °C for 3 hours before fixation in formalin for at 37 °C for 10 mins. Fixed platelets were added to a 24-well plate containing poly-L-lysine coated glass coverslips and centrifuged to immobilise platelets before washing and staining as described below.

### Immuno-labelling

Following PBS washes cells were permeabilised with 0.1 % Triton X-100 for 5 min and then washed in PBS and blocked for 30 min in block buffer (1 % BSA, 2 % goat serum in PBS). Tubulin was immuno-labelled with 2 *µ*g ml^-1^ anti-*α*-tubulin (Clone DM1A Sigma-Aldrich) diluted in block buffer for 1 h at room temperature. Tubulin was secondary labelled with 4 *µ*g ml^-1^ antimouse-Alexa647 and F-actin was labelled with 13 nM phalloidin-Alexa488 (Thermo-Fisher). Cells were washed and mounted in hydromount (National Diagnostics) prior to imaging.

### SDS-PAGE and western blotting

Lysates from platelets incubated for 3 hours with 0, 2 or 5 *µ*M SMIFH2 were prepared from platelets at 5 × 10^8^ ml^-1^ in an equal volume of 2x lysis buffer on ice for 10 minutes. Lysates were mixed with sample buffer and boiled for 5 minutes before cooling. Samples and protein marker were loaded into wells of pre-cast polyacrylamide gels (Bolt, Invitrogen) at 60 V for 15 minutes to stack proteins, followed by 120 V for 1 hour for separation. Proteins were transferred to PVDF membranes for 10 minutes using the Trans-Blot Turbo Transfer System (Bio-rad), then incubated for 1 hour at room temperature in blocking buffer (5 % BSA in TBST, filtered). Blots were incubated with primary antibodies against either *α*-tubulin (clone DM1A, Sigma-Aldrich), acetylated *α*-tubulin (Clone 6-11B-1, Cell signalling Technology) or tyrosinated *α*-tubulin (Clone YL1/2, Merck Millipore) in blocking solution at 4 °C overnight. Blots were washed 3 times for 10 minutes each in TBST, then incubated for 1 hour with anti-mouse or anti-rat secondary antibody, diluted 1:10000 in TBST, before a further 3 washes. Using LI-COR Odyssey imaging system and Image Studio software, western blots were exposed for 2-10 minutes and band intensities were quantified.

### Platelet aggregation and secretion

Platelet aggregation and dense granule ATP secretion was monitored using 300 *µ*L of washed platelets at 2 × 10^8^ mL^-1^. Stimulation of platelets was performed in a Chrono-Log aggregometer (Chrono-Log, Havertown, PA, USA) with continuous stirring at 1200 rpm at 37 °C and ATP secretion was determined during aggregation using Chronolume reagent (Chrono-Log). Where inhibitors were used these were added to the platelets for 10 mins prior to addition of platelet agonists.

### Microscopy

Images were acquired using an Axio Observer 7 inverted epifluorescence microscope (Carl Zeiss) with Definite Focus 2 autofocus, 63x 1.4 NA oil immersion objective lens, Colibri 7 LED illumination source, Hammamatsu Flash 4 V2 sCMOS camera, Filter sets 38 and 50 for Alexa488 and Alexa647 respectively and DIC optics. LED power and exposure time were chosen as appropriate for each set of samples but kept the same within each experiment. Using Zen 2.3 Pro software, five image stacks were taken per coverslip for fixed platelet experiments. For live cell experiments, images were taken at 23°C for 30 minutes at 5 second intervals using differential interference contrast (DIC) to visualise the extent of platelet spreading and Zeiss triple filter set 105 in combination with 488nm and 647nm LEDs to visualise LifeAct-GFP and SiR-tubulin. For resting human platelet experiments, confocal images were acquired using an SP2 confocal microscope (Leica Microsystems) with a 63x 1.4NA oil immersion objective lens and Ar 488 and HeNe 633 lasers. Five image stacks were taken per coverslip using Leica confocal software.

### Image analysis

Post capture image analysis was performed using Fiji ^[14]^. For fixed cell measurements, data are from means from three independent experiments, with between 150 and 300 individual platelets analysed for each treatment per experiment. For live cell spreading analysis, individual platelets were cropped from the field of view to allow synchronisation of spreading analysis. The spread platelet area was measured using ROI manager and Measure functions of Fiji. For live cell measurements data are from means from three independent experiments, with 5 platelets measured for each treatment per experiment.

### Statistics

Data analysis was carried out using Microsoft Excel and Graphpad Prism V6. Results are shown either as mean ± SEM or mean ± SD (as indicated in the figure legends) and statistical significance was analysed using One ANOVA and Dunnetts multiple comparison tests. P values are given in the text whilst in figures is represented by stars (P<0.05 = *; P<0.01 = **; P<0.005 = ***).

## Results

### Effect of global formin inhibition on platelet function

We had previously hypothesised that the lack of a phenotype in the mDia1 KO mice was due to a redundancy with other formin proteins expressed in platelets ^[8]^. Therefore we tested this by treating both mouse and human platelets with an FH2 domain inhibitor (SMIFH2) to establish if FH2 activity was required for proper platelet function.

### Mouse platelets

In mouse platelets treated with increasing concentrations of SMIFH2 and spread on fibrinogen, platelet spreading is significantly reduced (Fig. 1a). Control samples show that over 80 % of platelets display a nodule/filopodia phenotype, characteristic of mouse platelets spreading on fibrinogen. With increasing concentrations of SMIFH2 this deceases so that at 5 *µ*M approx. 90 % of platelets are unspread (Fig. 1b). Accordingly, platelet surface area was significantly reduced at both 4 *µ*M (P=0.027) and 5 *µ*M (P=0.013) SMIFH2 (Fig. 1c). Staining of platelet F-actin with Alexa488-phalloidin reveals that SMIFH2 causes disruption to F-actin organisation with actin forming spots in the cell and diffuse rings at the cell periphery (Fig. 1a). Tubulin organisation is also disrupted; at 5 *µ*M SMIFH2 platelets show neither microtubule coils, nor spread cell microtubule networks and at 5 *µ*M SMIFH2 the tubulin ring usually observed in control unspread platelets appears diffuse and the platelets are smaller in size (Fig. 1a).

**Figure 1.**
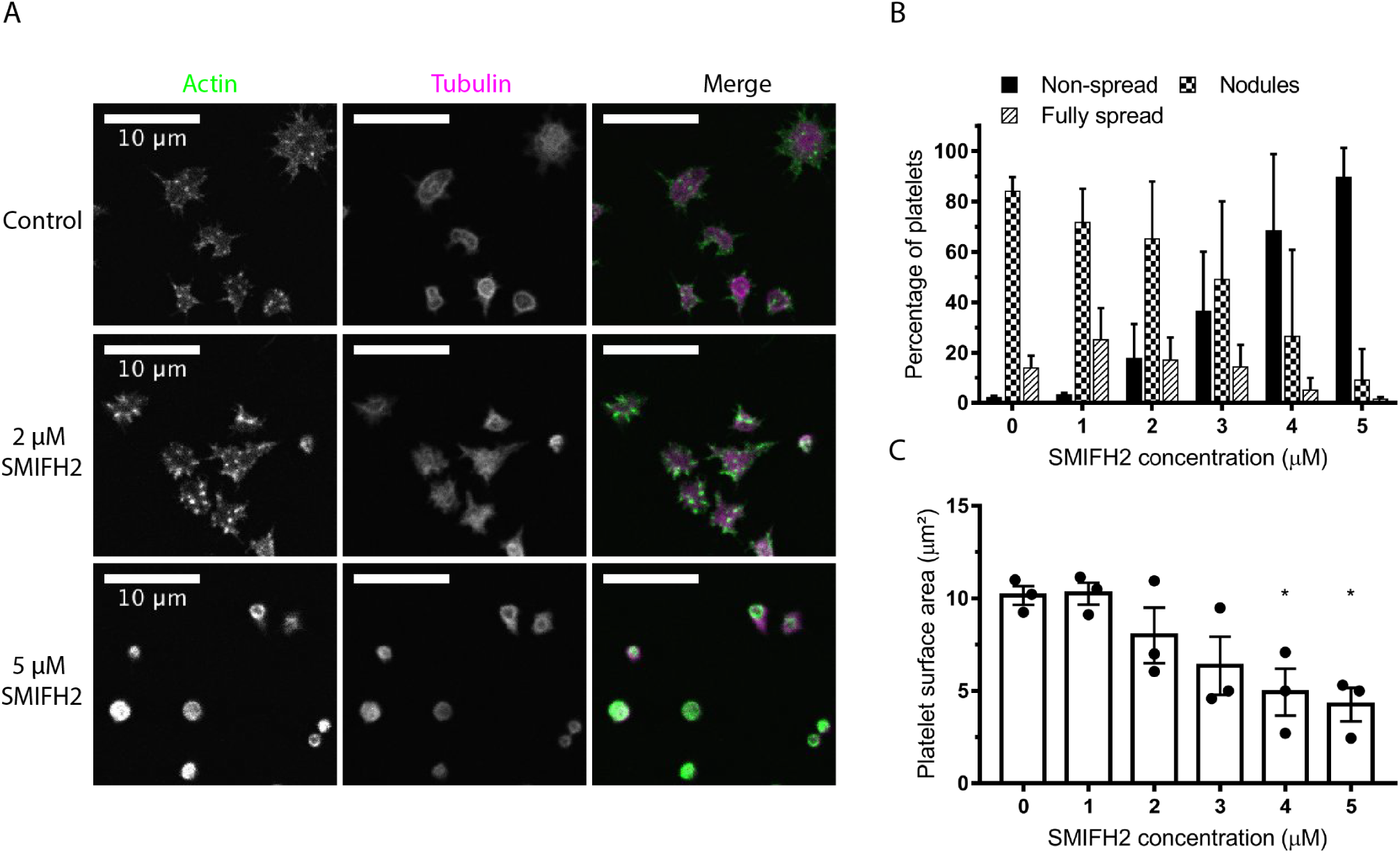
Inhibition of FH2 domains blocks mouse platelet spreading. **A)** Representative images of mouse platelets pre-treated with increasing concentrations of SMIFH2 prior to spreading on fibrinogen for 45 min. **B)** Analysis of spread platelet morphology (mean ± SD) and **C)** spread cell surface area (mean ± SEM) indicate a dose dependent inhibition of platelet spreading with almost complete spreading blockade at 5 *µ*M SMIFH2. Scale bars = 10 *µ*m.

### Human platelets

A similar effect on spreading is seen in human platelets to that observed in mouse platelets in that spreading is reduced (Fig. 2a). Control samples show that approx. 80 % of platelets display lamellipodia, stress fibres or are fully spread with an additional ≈ 15 % being at the nodule/filopodia stage. With increasing concentrations of SMIFH2, this decreases so that at 5 *µ*M SMIFH2 approx 90 % of cells are unspread with the remainder displaying occasional small filopodia or poorly formed lamellipodia (Fig. 2b). Platelet surface area was significantly reduced at 2 *µ*M (P=0.012), 3 *µ*M (P=0.003), 4 *µ*M (P=0.003) and 5 *µ*M (P=0.001) SMIFH2 (Fig. 2c). F-actin organisation is disrupted and at 5 *µ*M SMIFH2 no visible actin structures are present and the actin forms bright spots and with occasional diffuse rings at the cell periphery (Fig. 2a). Tubulin organisation is also disrupted and at 5 *µ*M SMIFH2 rings of tubulin are observed around the edge of the cell, although these appear different to the normal microtubule coils observed in unspread platelets. We also tested the effect of SMIFH2 on platelets spreading on fibrinogen and stimulated with 0.1 U ml^-1^ thrombin. Again, spreading is completely inhibited by 5 *µ*M SMIFH2 (surface area P=0.0001) (Supp. Fig.1). To test that the observed effects were not due to non-specific inhibition of platelet function, human platelets pre-treated with SMIFH2 were tested for aggregation and secretion responses to collagen and thrombin. No significant difference was observed for platelets treated with 5 *µ*M SMIFH2 (Supp. Fig. 1) indicating that the observed effects on spreading are not due to non-specific inhibition on platelet signalling.

**Figure 2.**
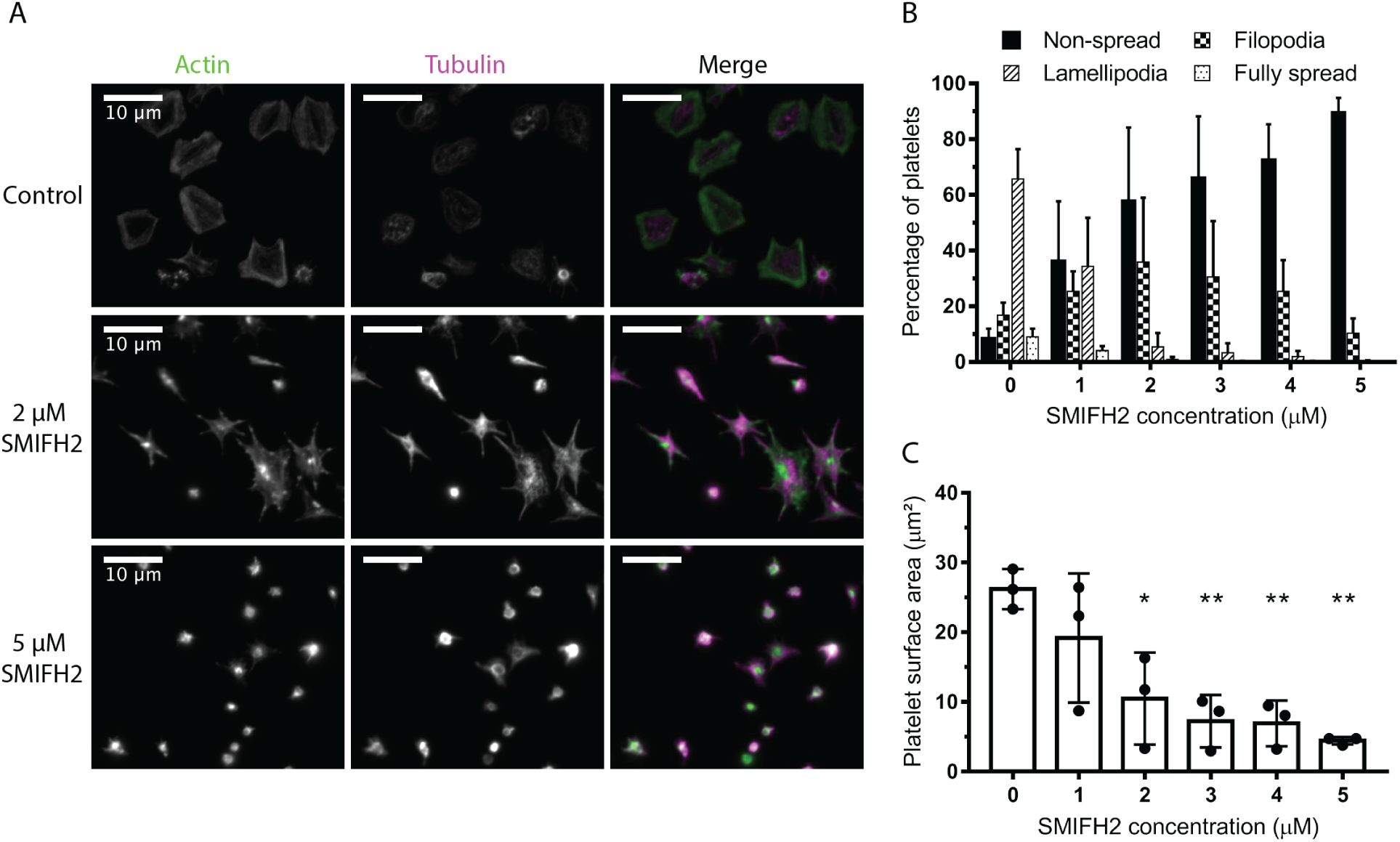
Inhibition of FH2 domains blocks human platelet spreading. **A)** Representative images of human platelets pre-treated with increasing concentrations of SMIFH2 prior to spreading on fibrinogen for 45 min. **B)** Analysis of spread platelet morphology (mean ± SD) and **C)** spread cell surface area (mean ± SEM) indicate a dose dependent inhibition of platelet spreading with almost complete spreading blockade at 5 *µ*M SMIFH2. Scale bars = 10 *µ*m.

### Live platelet spreading

To determine the effect of FH2 inhibition on cytoskeletal and spreading dynamics, platelet spreading on fibrinogen was followed in real time using DIC microscopy for both human and mouse platelets and spread platelet area measured at a range of time points. In control mouse platelets mean surfaces area remained fairly constant for the first 60 seconds as the platelets began spreading and extended and retracted filopodia. From about 60 seconds onwards, the platelets started to form lamellipodia and increased in size (Fig. 3a). A similar pattern was observed for control human platelets, although the formation of lamellipodia happened faster than in mouse platelets (Fig. 3b). In the presence of 5 *µ*M SMIFH2, both mouse and human platelets failed to spread and remained in this state for the duration of the imaging (Fig. 3a & b). For platelets treated with a lower concentration of SMIFH2, the spreading was reduced compared to controls (Fig. 3a & b) indicating that 2 *µ*M SMIFH2, there is still enough FH2 mediated actin polymerisation for limited platelet spreading.

**Figure 3.**
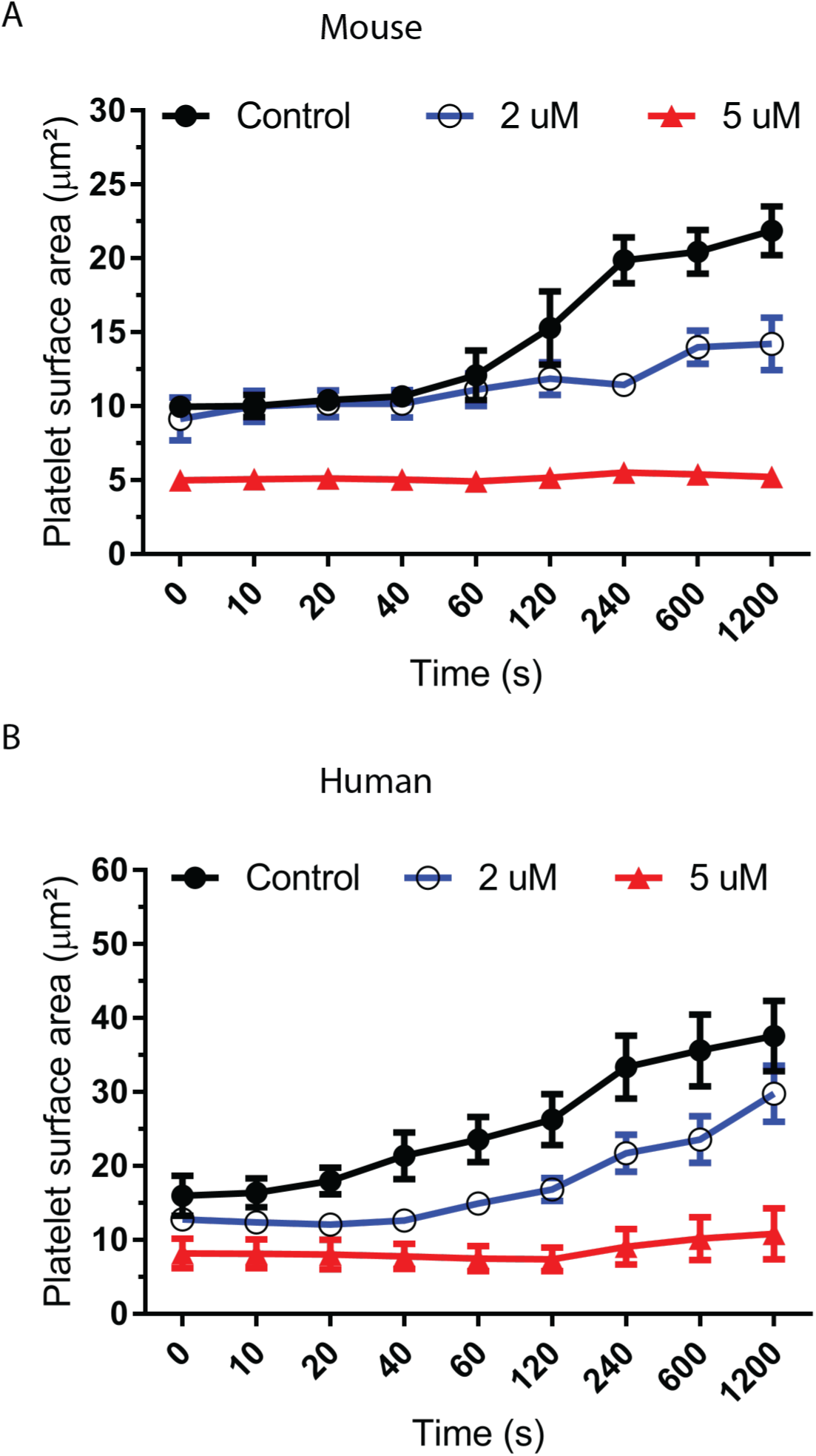
Inhibition of FH2 domains reduces spread area of live platelets. Live spreading analysis of **A)** mouse and **B)** human platelets on fibrinogen ± 2 or 5 *µ*M SMIFH2. (mean ± SEM)

To investigate this effect in more detail, platelets from the Lifeact-GFP mouse ^[13]^ were loaded with SiR-Tubulin ^[15]^ to allow visualisation of the F-actin and microtubule dynamics. Platelets were then allowed to spread on fibringen coated coverslips in the presence or absence of 5 *µ*M SMIFH2. Still images of representative platelets can be found in Fig. 4a (Control) and Fig. 4b (5 *µ*M SMIFH2). Example videos of can be found in Supplementary videos 1-4. In control cells, the characteristic spreading stages are observed including the generation of filopodia, actin nodules, lamellipodia and stress fibres as well as the twisting and contraction of the microtubule coil (Fig. 4a).

**Figure 4.**
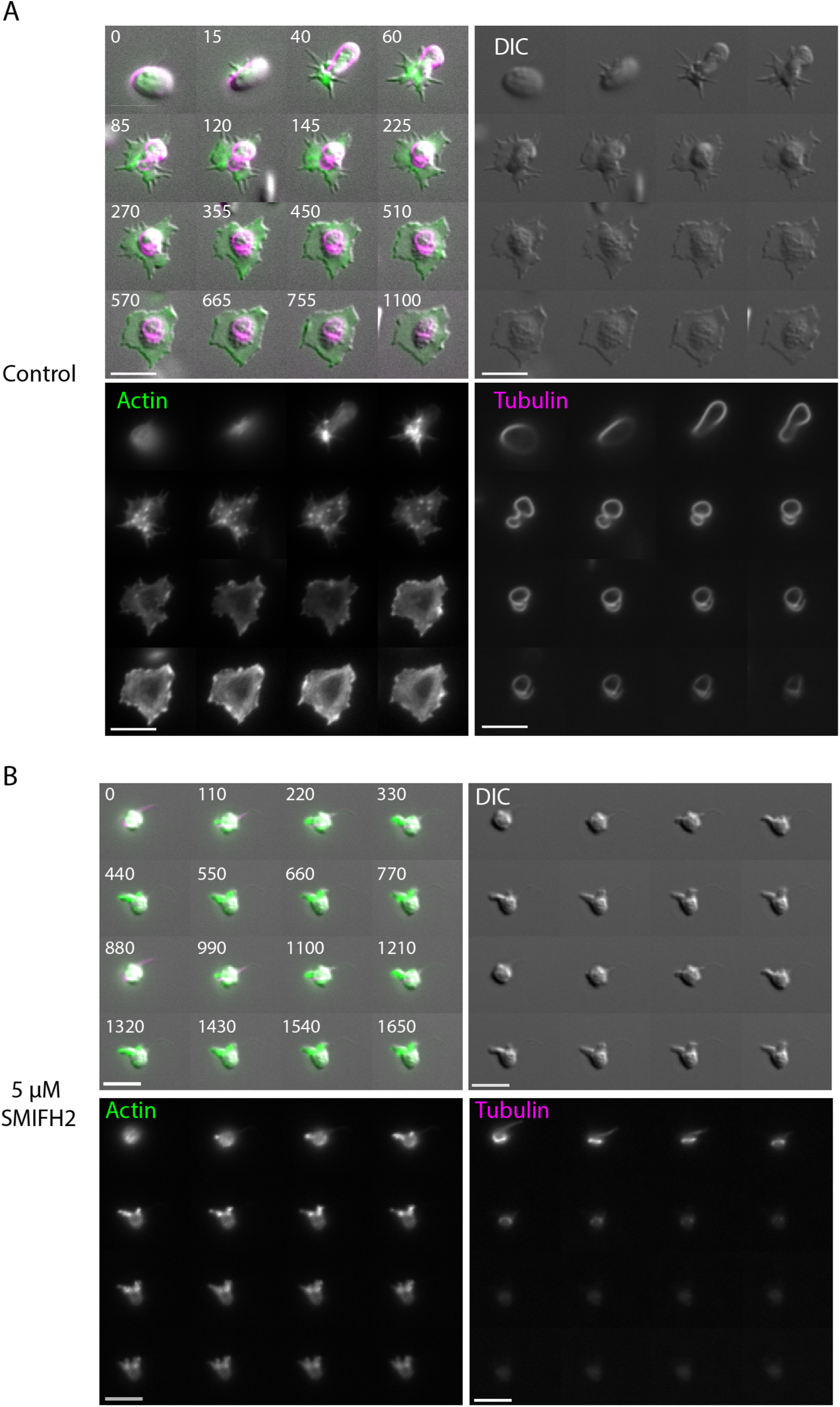
Inhibition of FH2 domains disrupts proper cytoskeletal organisation in spreading platelets. Representative timelapse images from **A)** control and **B)** 5*µ*M SMIFH2 treated Lifeact-GFP mouse platelets spreading on fibrinogen showing effect of FH2 inhibition on morphology, F-actin organisation and tubulin ring organisation. Scale bar = 5 *µ*m. Time in merged images is in seconds. (See also supplementary videos 1 - 4).

In platelets pre-treated with 5 *µ*M SMIFH2, spreading dynamics and cytoskeletal organisation are completely disrupted. Occasional small protrusions can be observed in treated platelets, however, these seemed to form by blebbing rather than by the conventional actin mediated extension. No evidence of ’normal’ filopodia or lamellipodia formation are observed, actin nodules appear, but are fewer in number and have much longer lifetimes than in control samples and the platelets do not form stress fibres. Tubulin dynamics were also disrupted in SMIFH2 treated cells; tubulin rings appeared smaller and thicker than in controls. The ring was also less dynamic than in controls and the characteristic twisting and coiling of the ring was not observed. The tubulin ring also seemed to depolymerise much quicker than in control cells which often displayed a remnant ring even when fully spread (Fig. 4b). Taken together this data shows that global inhibition of FH2 domains in mouse and human platelets disrupts actin and tubulin dynamics and prevents platelet spreading.

### Effect of formin inhibition on the resting platelet cytoskeleton

Human platelets with a constitutively active mDia1 ^[10]^ demonstrate a macrothrombocytopenia. To establish if the FH2 domain of formin proteins plays a role in this size increase, resting mouse and human platelets were treated with SMIFH2 before fixation and staining for the microtubule coil. Inhibition of FH2 domains in both mouse (Fig. 5a & b) and human (Fig. 5e & f) platelets caused a significant, dose dependent decrease in the surface area of resting platelets (Mouse - 2 *µ*M, P=0.007; 5 *µ*M, P=0.003. Human - 2 *µ*M, P=0.03; 5 *µ*M, P=0.002). However, in both cases the reduction in size is not caused by depolymerisation of the microtubule coil; SMIFH2 treated platelets still display an intact coil, but one which is significantly smaller in diameter and thicker, than in control cells (Mouse - 5 *µ*M, P=0.009. Human - 2 *µ*M, P=0.003; 5 *µ*M, P=0.0002) (Fig. 5c, d, g & h). This data therefore indicates that the FH2 domains of formin proteins are important for the maintenance of resting platelet size.

**Figure 5.**
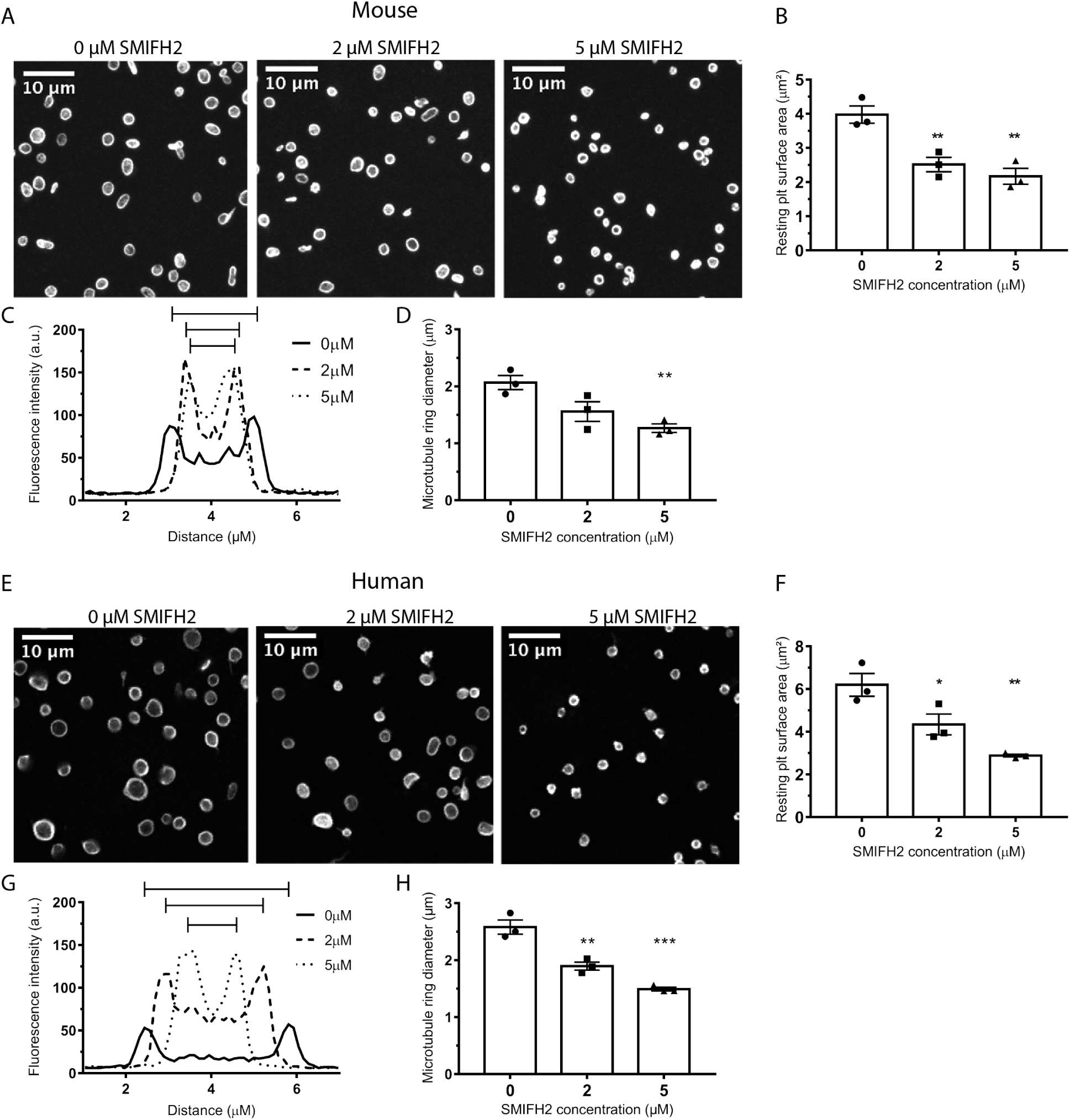
Inhibition of FH2 domains reduces resting platelet size. Representative images of **A)** mouse and **E)** human resting platelets incubated with SMIFH2 for 3 hours. Scale bars = 10 *µ*m. Analysis of resting platelet surface area indicates a significant decrease in platelet size upon treatment with SMIFH2 in both **B)** mouse and **F)** human platelets. Analysis of the microtubule rings of these platelets indicates that the microtubule ring remains intact upon SMIFH2 treatment, but their average diameter is reduced in both mouse **(C & D)** and human **(G & H)** platelets. For all plots data are mean ± SEM.

### Effect of FH2 inhibition on microtubule post-translational modifications

Formins can bind to microtubules via their FH2 domain and have been shown to regulate microtubule dynamics independently of effects on actin dynamics ^[16]^. Formins have been implicated in the acetylation of microtubules, a modification which, in combination with detyrosination, marks stable microtubules ^[17]^. We therefore tested the effect of formin FH2 domain inhibition on microtubule post-translational modification (PTM) in platelets. In resting human platelets treated with SMIFH2, a reduction was observed in the acetylation status and a small increase in the tyrosination status of *α*-tubulin (Fig. 6a). This resulted in a decrease in the acetylation:tyrosination status of tubulin in these platelets (Fig. 6b) which was significant at 5 *µ*M SMIFH2 (5 *µ*M, P=0.017). This data indicates that PTMs of *α*-tubulin in platelets is regulated, at least in part, by the action of the formin FH2 domain.

**Figure 6.**
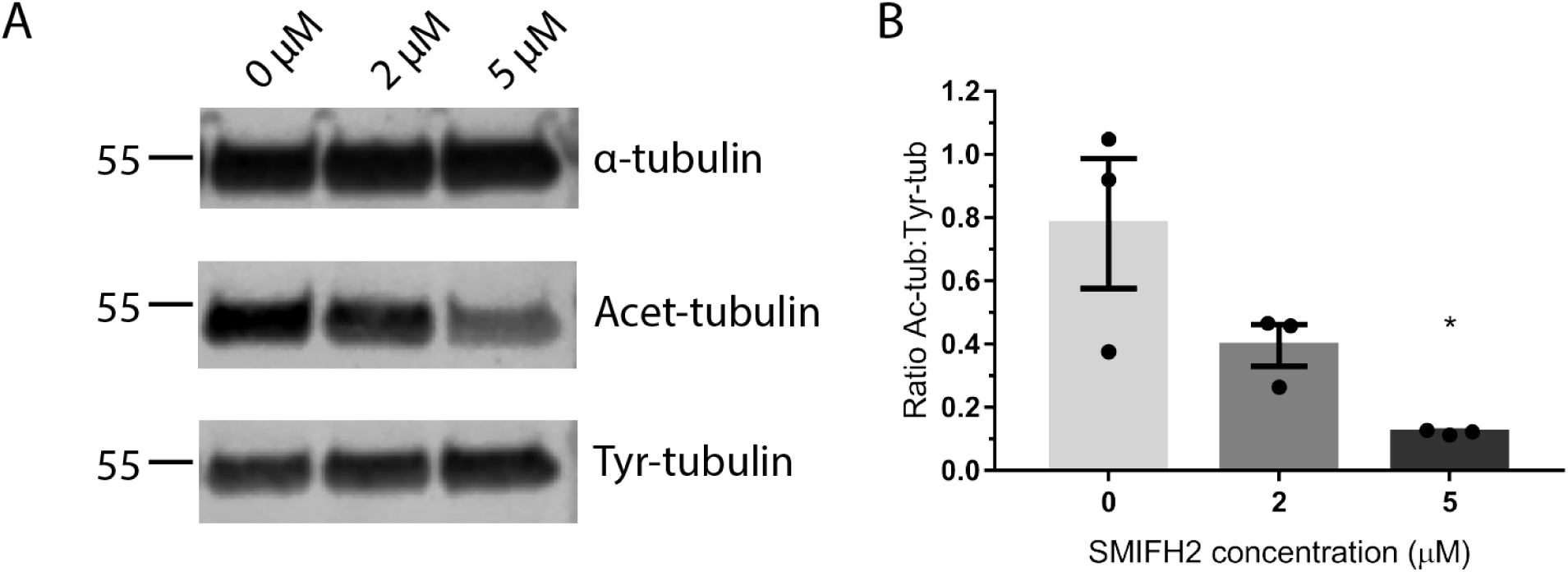
Inhibition of FH2 domains affects microtubule stability. **A)** Western blots of resting human platelet lysates probed for *α*-tubulin and its post-translational modifications, acetylation and tyrosination. **B)** Quantification of band intensities shows a reduction in the ratio of acetylated:tyrosinated tubulin in SMIFH2 treated cells (mean ± SEM).

## Discussion

The data presented in these studies indicates that the FH2 domain of formin proteins is required for normal cytoskeletal organisation and dynamics of platelets, as a dose dependent inhibition of platelet spreading by SMIFH2 was observed in both human and mouse platelets. It is well established that the FH2 domain is highly conserved across species and is absolutely required for the actin polymerisation action of formin proteins ^[4]^. Thus, the dose dependent effects of SMIFH2 on platelet spreading can be largely explained as a result of decreased actin polymerisation. The possible off target and cytotoxic effects of SMIFH2 have been investigated. The concentrations of SMIFH2 used here are lower and were for a for shorter time period than those used in many studies ^[12]^ and are well below the concentrations reported to have effects on p53 function and golgi organisation ^[18]^. Furthermore, platelets treated with 5 *µ*M SMIFH2 retain normal aggregation and secretion response indicating that the platelet are viable. Therefore, it is reasonable to conclude that formin mediated actin polymerisation is a key component of platelet spreading responses.

In addition to the disruption of platelet actin dynamics, changes in microtubule organisation and dynamics was also observed. Microtubule coils in platelets at early stages of spreading were disrupted and spread platelet microtubule networks failed to form. Furthermore, the dynamics of microtubules were affected with observed microtubule coils failing to undergo the characteristic twisting of control platelets (as observed by Diagouraga *et al*. ^[19]^) and appearing to depolymerise more rapidly than in controls. Thus it would appear that there is an effect of SMIFH2 on microtubules in addition to the inhibition of actin polymerisation. There is substantial evidence that formins can regulate microtubule dynamics, both via direct binding of formins to tubulin, as well as via interaction with microtubule associated proteins (reviewed by ^[7]^). However, whether the effects observed here on spread platelet microtubule dynamics are due to blocking a direct interaction of FH2/formins with microtubules, or through indirect effects on actin polymerisation is unclear. There is evidence that formins can stabilize microtubules independently of their effect on actin ^[16]^ and that competition for FH2 domains between actin filaments and microtubules regulates actin and microtubule interactions ^[20]^. Regardless of the mechanism, such studies highlight the need for orchestration of the actin and microtubule cytoskeletons for proper platelet function which is clearly lacking in platelets treated with SMIFH2.

The importance of this cytoskeletal orchestration is supported by a key observation in this work, namely the effect of FH2 domain inhibition on the resting platelet. It is well characterised that depolymerisation of the microtubule coil, as in the case of treatment with microtubule deploymerising agents or by chilling of platelets, results in a reduced platelet size. However, here the decrease in size was not accompanied by a microtubule depolymerisation and rather the microtubule coil was intact, but smaller in diameter and appeared thicker in size, as if it had coiled in on itself. To establish a possible mechanism by which this might occur, we looked at the PTM status of tubulin in resting platelets when incubated with SMIFH2, and determined that the ratio of acetylated:tyrosinated tubulin is reduced when FH2 domains are inhibited. Thurston *et al*. ^[17]^ demonstrated that microtubule acetylation was a general feature of FH2 domains, but it remained unclear whether this is due to activation of a specific tubulin acetyltransferase (TAT) or inhibition of histone deacetylases (e.g. HDAC6) ^[21]^. Recent evidence suggests that formins play a dual role by stabilising microtubules to allow tubulin acetylation as well as by increasing the transcription of TAT1 ^[22, 23]^, although clearly the later will not be occurring in platelets treated with SMIFH2.

Tubulin PTMs have been shown to be important in regulating both microtubule stability and their interaction with microtubule associated proteins and are part of a tubulin code that regulates cytoskeletal dynamics ^[21, 24, 25]^. The changes reported here could affect the platelet microtubule coil in a number of ways; i) tubulin acetylation is associated with the protection of microtubules from mechanical stress ^[25]^ & ii) increased tyrosination is associated with increased kinesin motor activity ^[26]^. Together, these may explain the changes in microtubules observed in resting platelets upon FH2 inhibition. Increased kinesin motor activity combined with a reduction in microtubule – actinomyosin crosslinking causes the microtubule coil to decrease in size, reducing the size of the resting platelets. In addition decreased acetylation of microtubules makes them more prone to mechanical stress and may be why the microtubule coil depolymerises quicker in spreading platelets treated with SMIFH2.

There is a precedent in the literature regarding formin proteins, platelets and microtubule PTM status. Both Pan *et al*. and Stritt *et al*., have reported altered microtubule PTM in models of platelet and megakaryocyte formin disruption ^[9, 10]^. Although the genetic approaches employed in those studies resulted in a loss or reduction of protein expression, we have demonstrated here that blocking the action of platelet formin FH2 domains alone is sufficient to disrupt microtubule PTMs and subsequent function.

In conclusion, we demonstrate that the FH2 domain of formin proteins plays a key role in the organisation and dynamics of the platelet actin and tubulin cytoskeletons. The fact that mDia1 knockout platelets show no spreading phenotype, would seem to support the hypothesis that multiple formin proteins can fulfill this role during platelet activation and thus the challenge is now to identify the contribution that each of the platelet expressed formin proteins makes to this process.

## Acknowledgements

We thank the British Heart Foundation (PG/15/114/31945), University of Birmingham Medical School and COMPARE for funding. The authors would like to thank the Biomedical Services Unit (BMSU) for management of the Lifeact-GFP and WT mice colonies. We would also like to thank Professor Steve P Watson for his continued support and the camel for inspiration.

## Authorship Contributions

These contributions follow the Contributor Roles Taxonomy guidelines (CREDIT) which can be found at casrai.org/credit

**Conceptualization:** S.G.T.;

**Funding acquisition:** S.G.T.;

**Investigation:** M.Z., H.L.H.G., S.G.T.;

**Project administration:** S.G.T.;

**Supervision:** M.Z., S.G.T.;

**Visualization:** M.Z., H.L.H.G., S.G.T.;

**Writing** – **original draft:** M.Z., S.G.T.;

**Writing** – **review & editing:** M.Z., H.L.H.G., S.G.T.

## Conflict of interests

The authors declare no competing financial interests.

