## Supplementary figures and videos for "The FH2 domain of formin proteins is critical for platelet cytoskeletal dynamics"

---

### Supplementary Information

#### Supplementary Videos

#### Supplementary Figures

### Supplementary Videos and Legends

#### Supplementary Video 1

Example time-lapse of control treated mouse platelets spreading on fibrinogen and imaged for morphology, F-actin and microtubules.

#### Supplementary Video 2

Example time-lapse of control treated mouse platelets spreading on fibrinogen and imaged for morphology, F-actin and microtubules.

#### Supplementary Video 3

Example time-lapse of 5 $\mu$M SMIFH2 treated mouse platelets spreading on fibrinogen and imaged for morphology, F-actin and microtubules.

#### Supplementary Video 4

Example time-lapse of 5 $\mu$M SMIFH2 treated mouse platelets spreading on fibrinogen and imaged for morphology, F-actin and microtubules.

### Supplementary Figures

### Supplementary Figure Legends

**Supplementary Figure 1**.

Effect of SMIFH2 on **A)** platelet spreading (in the presence of 0.1 U/ml$^{-1}$ thrombin) **B)** aggregation and **C)** ATP secretion in response to collagen and thrombin.

**Supplementary Figure 2**.

Full western blots for data in Figure 6a.

**Supplementary Figure 3**.

Visual abstract for the manuscript.

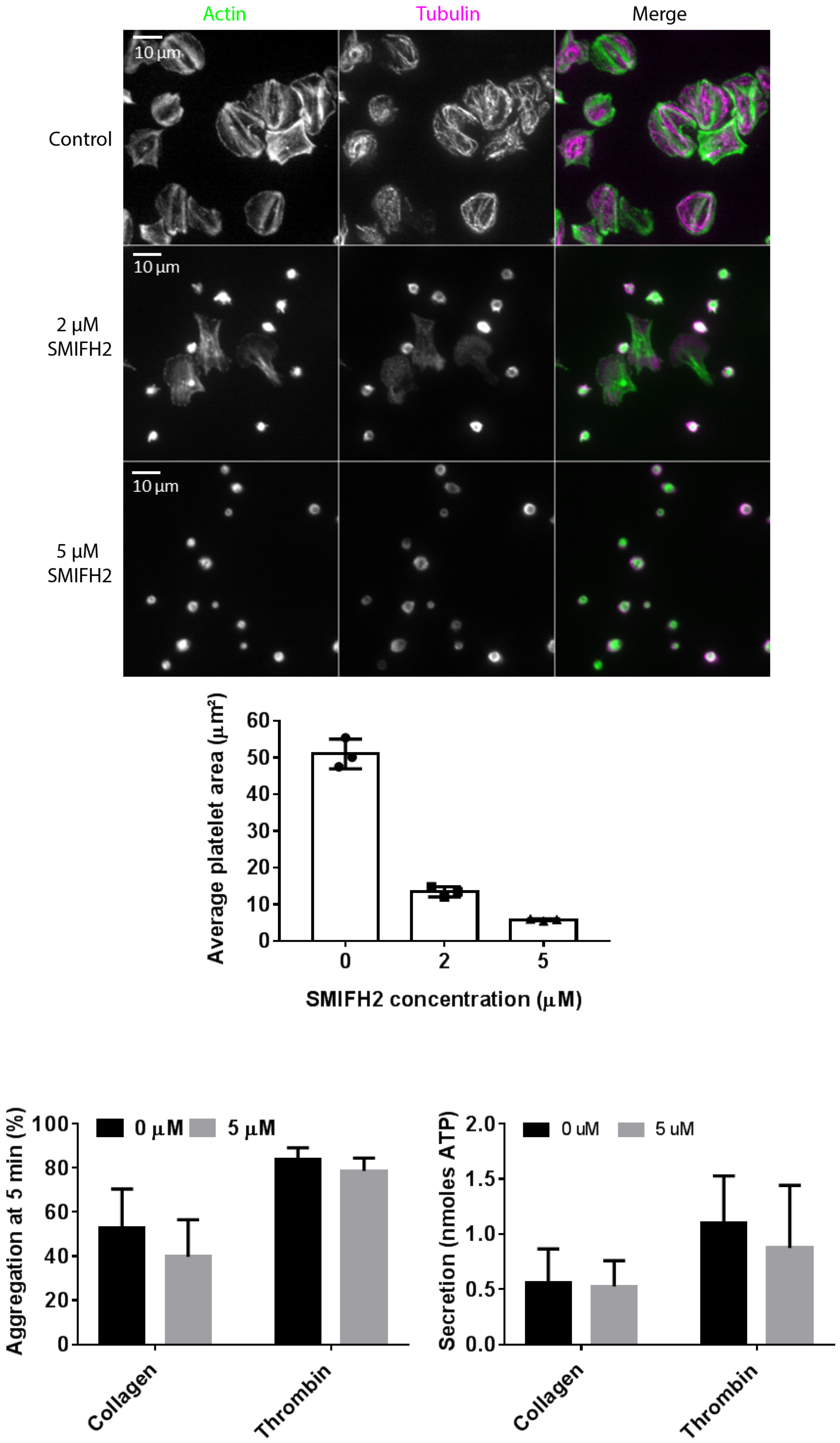
Actin
Tubulin
Merge
Control
2 µM SMIFH2
5 µM SMIFH2
10 µm
10 µm
10 µm
Average platelet area (µm²)
60
50
40
30
20
10
0
0
2
5
SMIFH2 concentration (µM)
Aggregation at 5 min (%)
100
80
60
40
20
0
0 µM
5 µM
Collagen
Thrombin
Secretion (nmoles ATP)
2.0
1.5
1.0
0.5
0.0
0 uM
5 uM
Collagen
Thrombin

Figure 1:

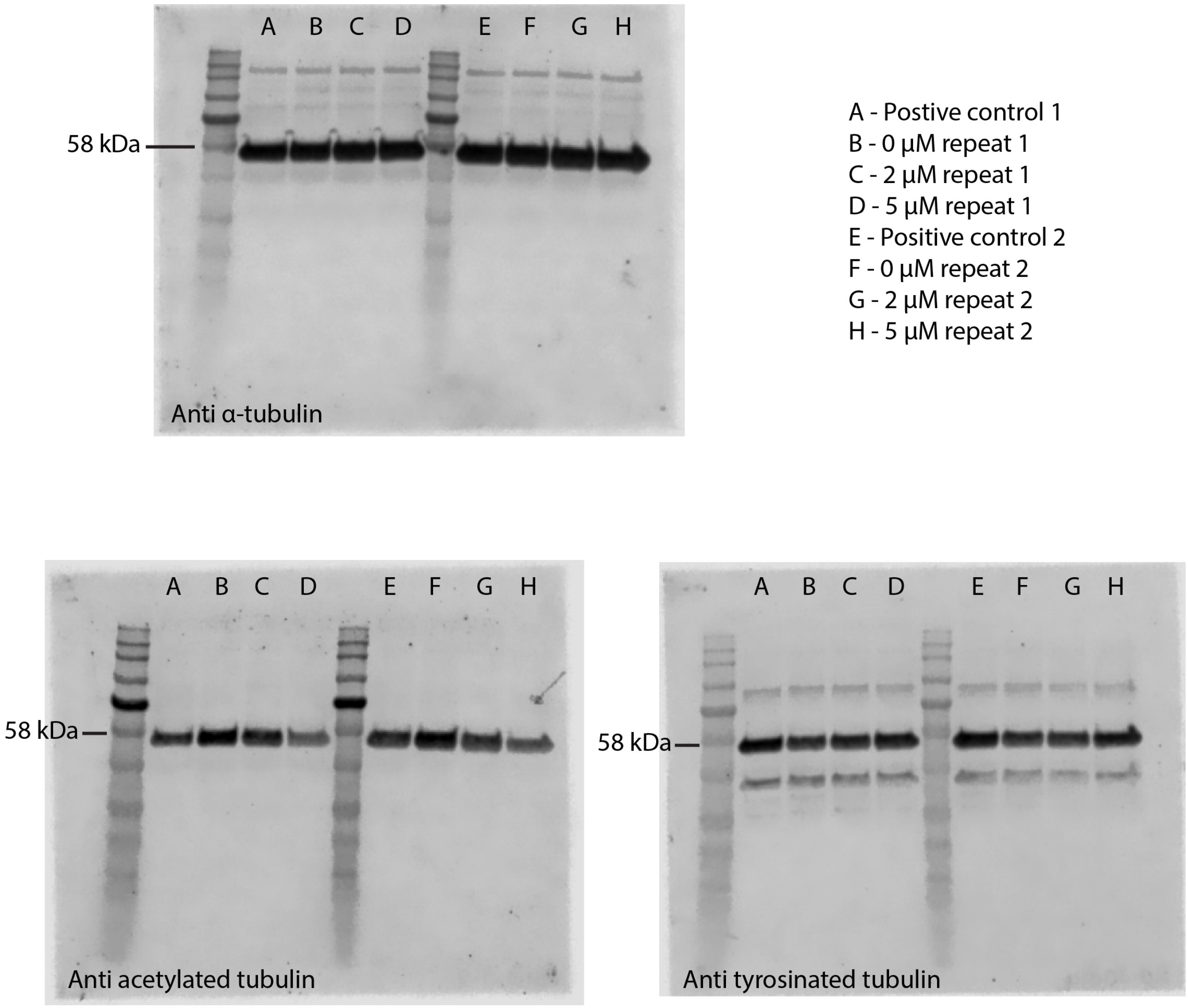
A B C D E F G H
58 kDa
Anti α-tubulin
A - Postive control 1
B - 0 μM repeat 1
C - 2 μM repeat 1
D - 5 μM repeat 1
E - Positive control 2
F - 0 μM repeat 2
G - 2 μM repeat 2
H - 5 μM repeat 2
A B C D E F G H
58 kDa
Anti acetylated tubulin
A B C D E F G H
58 kDa
Anti tyrosinated tubulin

Figure 2:

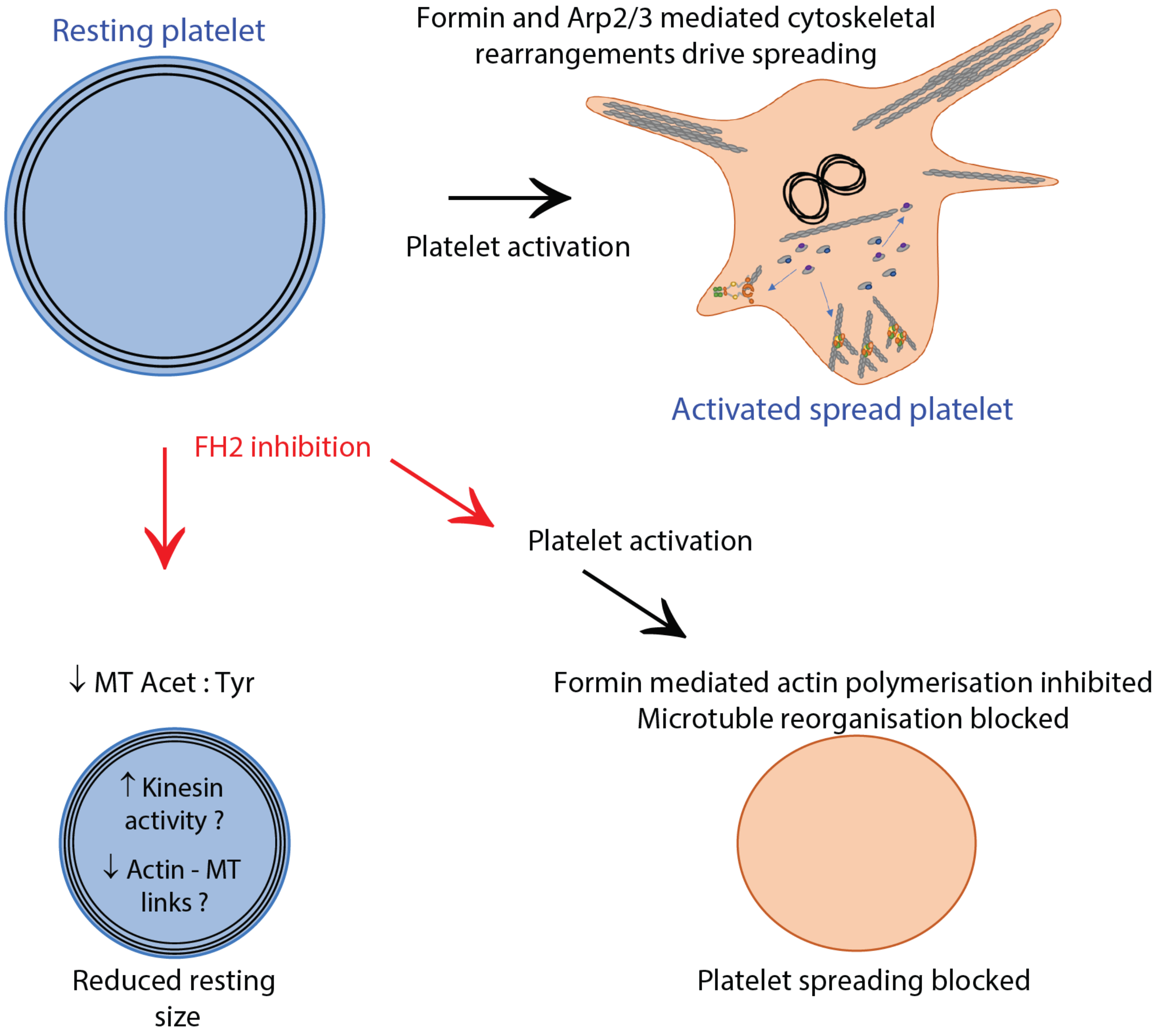
Resting platelet
Formin and Arp2/3 mediated cytoskeletal rearrangements drive spreading
Platelet activation
Activated spread platelet
FH2 inhibition
Platelet activation
↓ MT Acet : Tyr
↑ Kinesin activity ?
↓ Actin - MT links ?
Reduced resting size
Formin mediated actin polymerisation inhibited
Microtuble reorganisation blocked
Platelet spreading blocked

Figure 3:
